# Bile canaliculi contract autonomously by releasing calcium into hepatocytes via mechanosensitive calcium channel

**DOI:** 10.1101/449512

**Authors:** Kapish Gupta, Ng Inn Chuan, Binh P. Nguyen, Lisa Tucker-Kellogg, Boon Chuan Low, Hanry Yu

## Abstract

Bile canaliculi (BC) are the smallest vessels of the biliary tree. They are formed from the apical surfaces of adjoining hepatocytes, resulting in lumenal conduits for bile flow. Bile is propelled along the BC by hepatocyte contractions that arise from cyclic waves of apico-basal Ca^2+^, but the source and regulation of Ca^2+^ has been unclear. We report that BC contraction correlates with cyclic transfer of Ca^2+^ from BC lumen to apico-basal Ca^2+^ waves in adjacent hepatocytes, and does not correlate with endoplasmic reticulum Ca^2+^. BC contractility was triggered by ionophore A23187 and unaffected by Thapsigargin. The cycles of Ca^2+^ transfer could be blocked by the mechanosensitive calcium channel inhibitor GsMTx-4, resulting in cholestatic generation of BC-derived vesicles. The mechanosensitive calcium channel Piezo-1 is preferentially localized at BC membranes, and its hyper-activation by Yoda1 causes increased Ca^2+^ transfer and increased BC contractility. We propose that canaliculi achieve biomechanical homeostasis through the following feedback system: the pressure of accumulated bile is sensed by mechanosensitive channel, which transmit biliary calcium into adjacent hepatocytes for contraction of the BC lumen and propulsion of the bile.

## Introduction

Lumenal contractility plays an important role in multiple physiological processes such as fluid flow and bowl movement. In these systems of contractile propulsion, the trigger for contraction is the content itself. For instance, the food we eat triggers sensory neurons that regulate smooth muscle contraction along the gastrointestinal tract to drive bowl movement. Some finer lumenal structures lack muscle layer but exhibit peristaltic-like contraction driven by myosin. For instance, bile is secreted into a branched network of intra-hepatic lumen known as bile canaliculi (BC)^1^. Secretion and segregation of bile is important for liver physiology, as any breach can lead to jaundice or cholestasis^2^. BC show peristaltic motion to facilitate flow of bile contents into the bile duct^3^. Studies of BC dynamics are generally performed with *in vitro* culture systems,^4^ which have been shown to exhibit periodic expansion and contraction^5^, reminiscent of the *in-vivo* peristaltic motion^6^. BC expand due to activity of transporters on the BC membrane^7^; and BC contracts due to the actomyosin contractility triggered by increased intracellular Ca^2+^ ^8^ ^9^. Ca^2+^ micro-injection studies by Watanabe *et al.*^10^ and subsequent studies show that the increase in intracellular Ca^2+^ can induce BC contractions^11-14^. However, the source of Ca^2+^ remains elusive. One theory suggests that BC contraction is regulated by extracellular Ca^2+^ ^14, 15^, while another suggests that BC contraction is regulated by Ca^2+^ release from the endoplasmic reticulum (ER)^16^. In this study, we utilized various Ca^2+^ probes and live imaging of BC contractions to trace the source and propagation of Ca^2+^ waves associated with BC contractions. We found that BC contractions are triggered by the release of Ca^2+^ from the BC lumen, corresponding to a transient Ca^2+^ increase in the cytoplasm adjacent to the BC. We did not find any correlation between BC contractions and the release of Ca^2+^ from the ER. We therefore deduce that BC regulate their own contractions by providing Ca^2+^ to surrounding cells. Furthermore, we hypothesize that the BC membrane can sense canalicular pressure via mechanosensitive channel (MCC); and that BC membrane tension acts as a homeostatic switch leading to BC contractions.

## Result

### Ca^2+^ is concentrated in BC

Hepatocytes were isolated from rat using collagenase perfusion^17^ and grown in collagen sandwich cultures to support the maintenance of hepatocyte polarity. BC developed 24-48 hours after cell seeding^18^. The BC appear as bright tube-like structures^19^ along the cell-cell interface (Figure 1A). They are dynamic structures that expand and contract periodically (Supporting Video 1)^1, 5-7^, ^16^. BC contraction can be dependent or independent of BC-derived vesicles (BCV) (Figure 1B and C). In sandwich culture, in the presence or absence of up to 1% DMSO, 80-90% contractions occur via BCV-independent mechanism while 10-20% contractions happen via BCV formation. Here we will first focus on the BCV-independent contractions.

**Figure 1:**
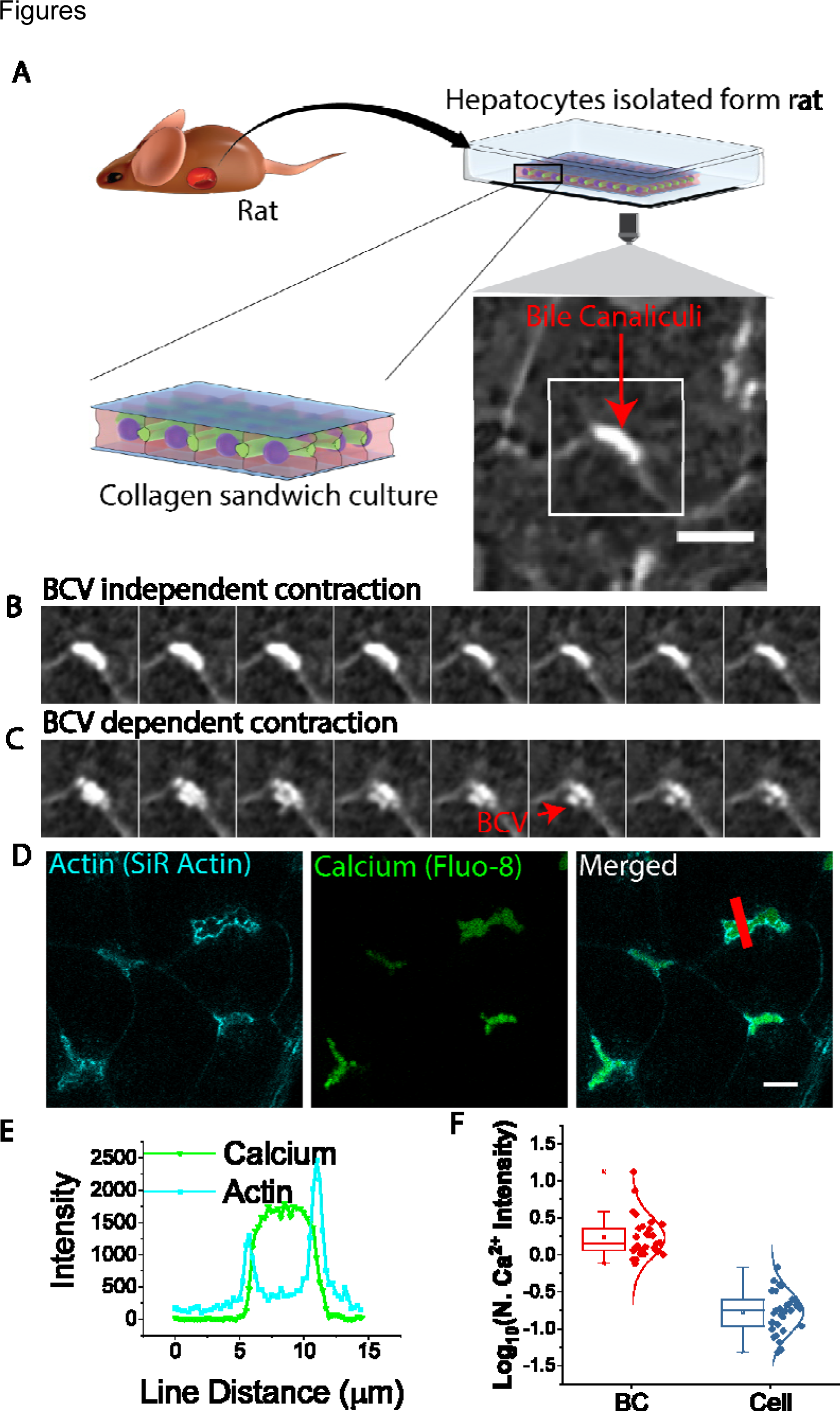
Ca^2+^ is present inside BC which can dynamically expand and contract. A) Schematic showing isolation of rat hepatocytes and culturing in collagen sandwich. Hepatocytes form tubular structure known as BC. These structures appear as bright tubes in phase contrast microscopy where BC actively expand and contract. A single BC can contract via one of the two ways. B) BC can contract via unknown mechanism of volume reduction possibly due to para-cellular leakage; C) BC can contract via BCV formation, which is the pinching of BC segment to reduce the BC volume. ∽85 % of contractions are driven by the first contraction mechanism and ∽ 15% via BCV formation. D) BC also store high concentration of Ca^2+^. BC are labelled for Actin (cyan from SiR actin), and for Ca^2+^ (green from Fluo-8). E) A line scan across a BC (red box in 1D) reveals that Ca^2+^ is present inside BC. F) Ca^2+^ intensity is much higher in BC than cells, suggesting that BC is the Ca^2+^ source. All scale Bar 10 µm

**Figure 2:**
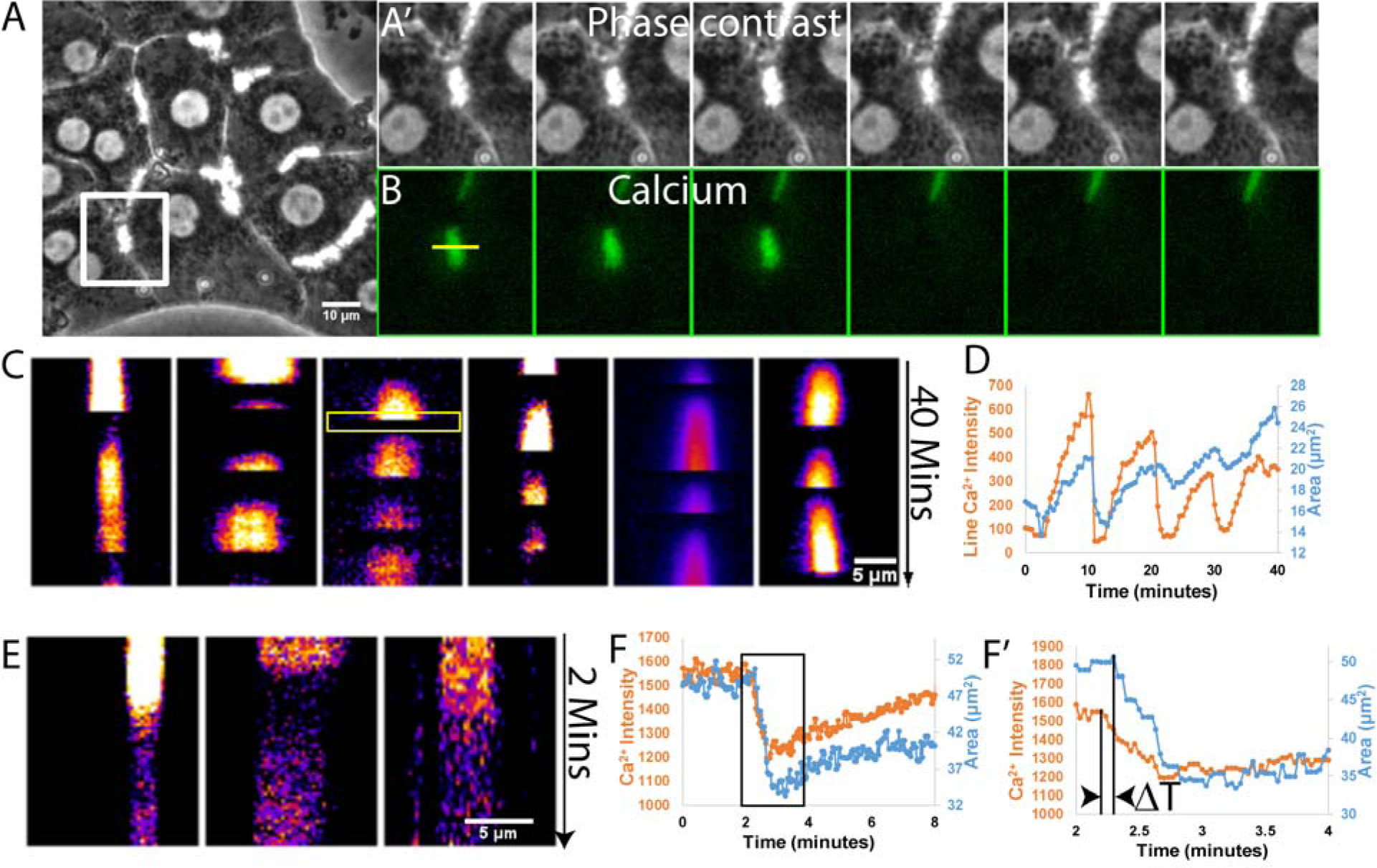
Ca^2+^ level fluctuates in BC and the BC Ca^2+^ level drops prior to BC contraction. A) Time-lapse images (30 second interval) of BC (A’) and Ca^2+^ (B) in cultured hepatocytes. Green signal disappearance (decrease in Ca^2+^ intensity) is associated with the decrease in BC area (BC contraction). C) shows Ca^2+^ intensity fluctuation as kymographs for 6 BC from 3 independent experiments. Third kymograph is drawn across yellow line in B and yellow box highlights kymograph for time lapse shown in B. D) Correlation of the BC area change and the Ca^2+^ intensity fluctuation. Ca^2+^ steadily increases in BC and then drops precipitously in periodic cycles always associated with BC contraction cycles. To investigate whether the BC Ca^2+^ drop triggers BC contraction in each cycle, we imaged BC labelled with Fluo-8 and SiR actin at high temporal resolution (1-3 sec interval). E) Kymograph showing fast Ca^2+^ intensity fluctuation at high temporal resolution. F) BC Ca^2+^ drops ∽10s prior to a BC contraction (F’ is the region highlighted by black box in F) supporting that BC Ca^2+^ drop triggers rather than the result of BC contraction.

BC contractions might be due to the Ca^2+^ release from the ER near BC^15^ or extracellular Ca^2+16^. However, the exact Ca^2+^ source regulating BC contraction has not been determined. We imaged the hepatocytes incubated with a Fluorescent Ca^2+^ marker (either Calcium Green™-1 AM (thermos fisher), Fluo-4 or Fluo-8) and found that Ca^2+^ is highly localized in the BC (supporting figure 1A). Fluo-8 was used as it was stable for long-term live cell imaging. A few earlier reports have shown localization of ER Ca^2+^) surrounding BC; however, our observation is the first report that Ca^2+^ is concentrated inside BC instead of localized on BC membrane where ER is present (Figure 1D). We used live actin dye (SiR actin, cytoskeleton) to label actin and imaged Z stack using the confocal microscope. After 3D reconstruction (supporting Video 2), we found that Ca^2^ is localized inside BC (Figure 1E and supporting figure 1A-B). The visualization of Ca^2+^ requires either wide-field imaging or confocal imaging at ∽2-4 µm above the cell-substrate contact since BC is lumenal and positions ∽1.5 µm above the cell-substrate contact plane. Most of the previous research on Ca^2+^ waves in hepatocytes was performed near the cell-substrate^15^ contact plane using either TIRF or confocal imaging; thus, BC Ca^2+^ was not observed in those studies.

The Ca^2+^ intensity was much higher in BC than in the cells (Figure 1F and supporting figure 1C). Due to extremely high BC Ca^2+^, it is difficult to image Ca^2+^ concurrently in both the ER (inside a cell) and BC (supporting figure 3D and E). We have adjusted the laser power and acquisition setting so as it is only sensitive to changes in BC Ca^2+^ and not to any changes in Ca^2+^ level in the cells. Presence of such a high concentration of Ca^2+^ inside BC intrigued us and we hypothesized that the canalicular Ca^2+^ may play a role in BC contractions.

**Figure 3:**
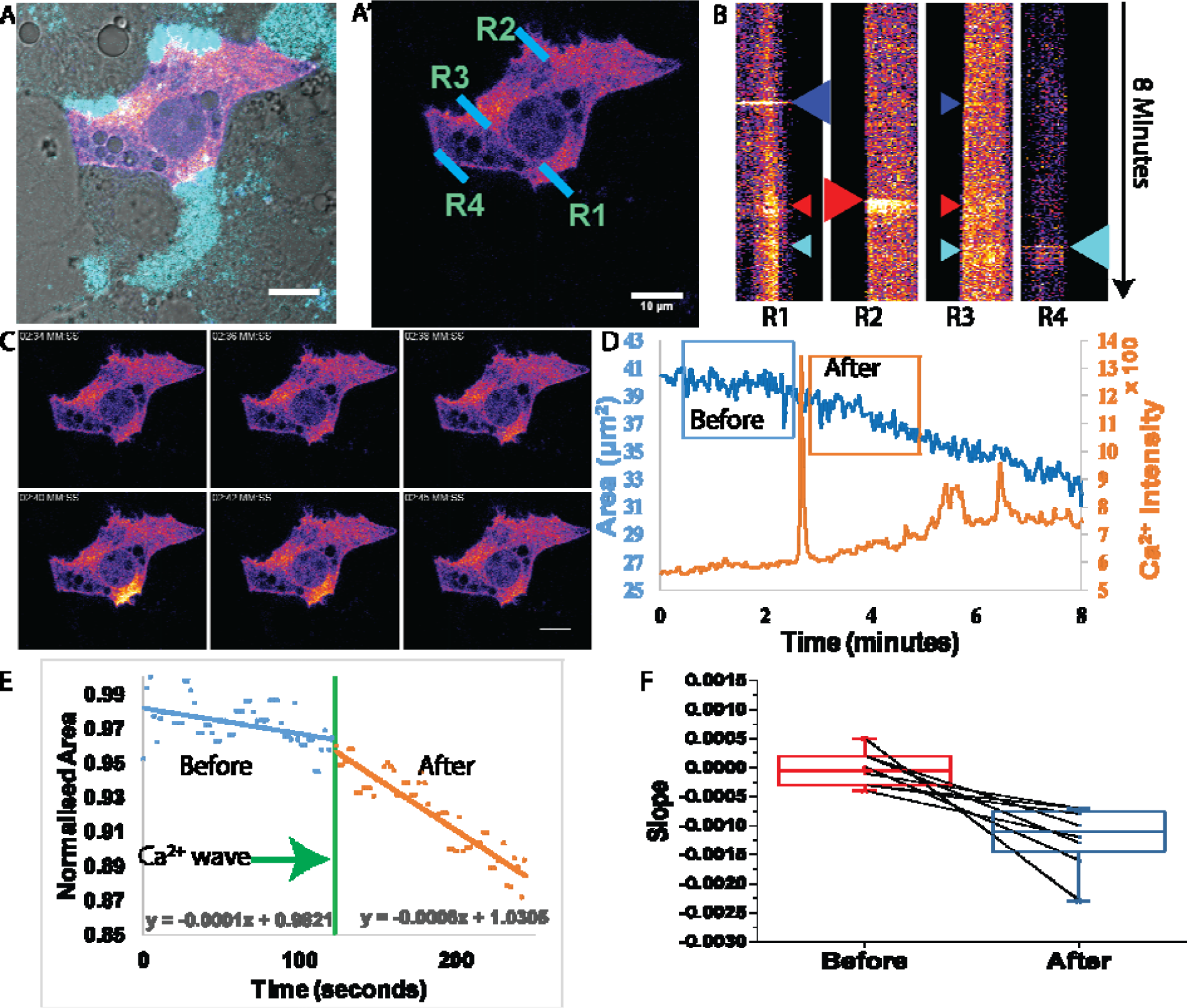
Apical Ca^2+^ triggers BC contraction. A) Shows a hepatocyte transfected with GCaMP (a Ca^2+^ indicator) and BC are marked in cyan. A’) shows the same cell with the only Ca^2+^ channel showing 4 different regions marked R1-R4. R1, R2 and R3 are along three distinct BC and R4 is along cell-cell contact. B) shows kymograph drawn across region R1-R4. As R1 shows transient increase in intensity (Large blue arrowhead), this increase in intensity around R1 is followed by slight increase at the other regions (small blue arrow). Similarly, the Ca^2+^ wave can occur at R2 (Large red arrow). This is followed by slight increase in Ca^2+^ intensity at R1 and R2 (small red arrow) suggesting travelling Ca^2+^ wave. Ca^2+^ waves can be generated via BC independent mechanism such as transport across cell membrane via gap junction or Ca^2+^ release from ER. One such example is shown by Ca^2+^ increase in R4 (large cyan arrow). This led to travelling Ca^2+^ wave (small cyan arrow), however not resulting in any contraction. Some BC (R3) do not show any Ca^2+^ wave during the imaged period and also did not show any contraction (Supporting figure 4C). C) A few time stamps (2.2 second interval) of the cell shown in Figure 3A showing Ca^2+^ wave at R1 as the Ca^2+^ intensity increases instantaneously at BC. D) shows the correlation of these Ca^2+^ waves with the BC contraction. BC area is steady before the Ca^2+^ peak (at ∽2.5 minute) and can be seen decreasing after Ca^2+^ wave. There are small Ca^2+^ peaks at around 5.5 and 6.5 minutes. Those were due to travelling wave generated at R2 and R4 (small red and cyan arrow in 3B). In this example low laser power was used and hence Ca^2+^ spikes around BC are much more visible than Ca^2+^ waves. Ca^2+^ waves are more visible at higher laser power as shown in supporting figure 4. E) shows the line fitting for area changes observed for two minutes before (blue box in D) and after (orange box in D) Ca^2+^ wave. There is a difference in slope before and after Ca^2+^ wave and slope become negative indicating BC contraction. E) Quantification of the slope for 8 such BC contractions. The slope was near ‘0’ prior to Ca^2+^ wave indicating no change in the BC area. Following Ca^2+^ wave all the slopes decrease indicating contraction. Ca^2+^ waves in hepatocytes can trigger BC contractions. All scale bar 10 µm

**Figure 4:**
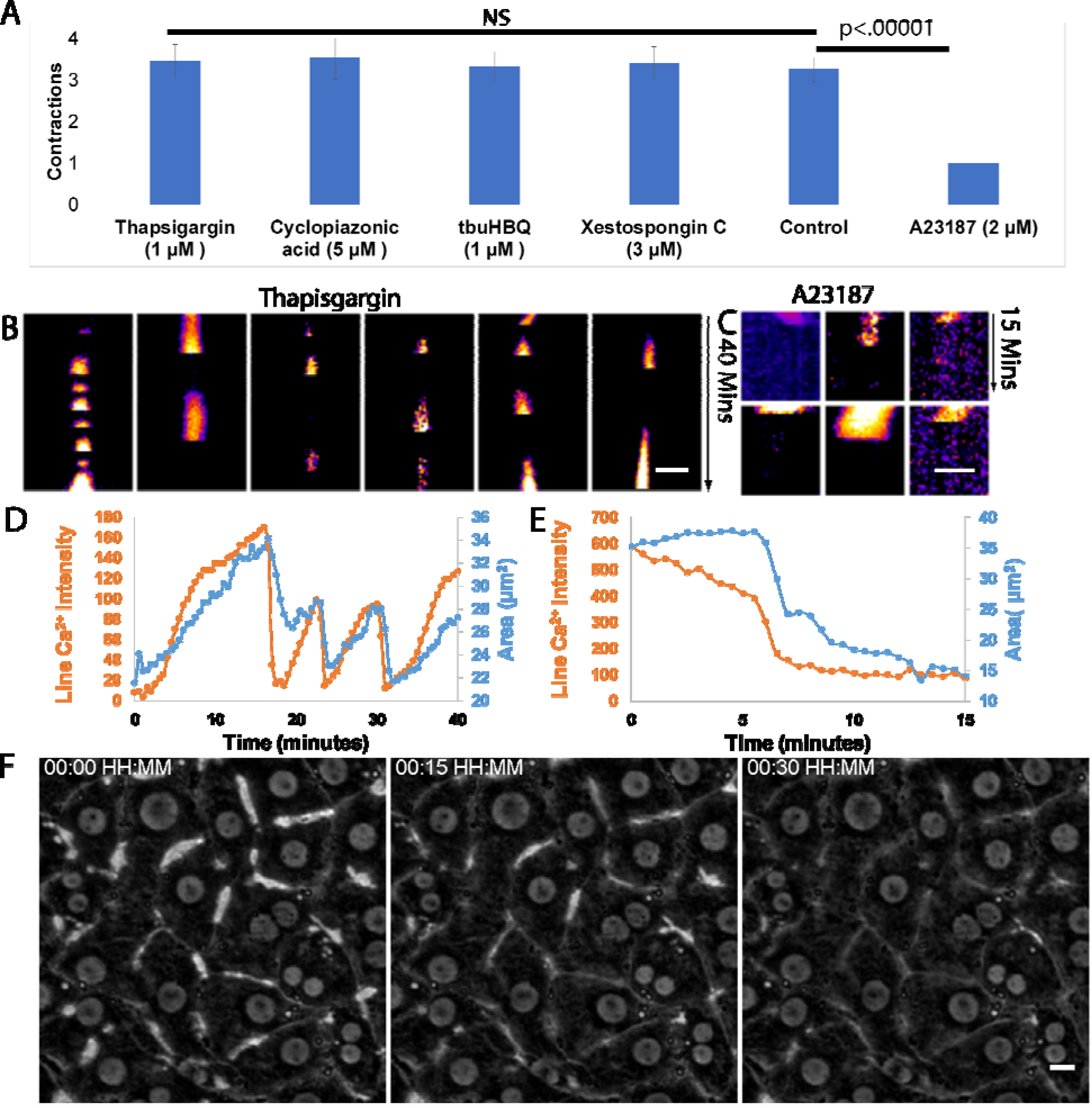
BC contraction and BC Ca^2+^ cycles are independent of the ER Ca^2+^. A) cumulative analysis of BC contractions using various ER and IP3 inhibitors showing no difference in the number of BC contractions observed in 40 minutes compared to a single contraction observed for A23187. BC Ca^2+^ cycles are maintained in presence of 1 µM Thapsigargin, a specific and potent inhibitor of SERCA. C) BC contraction are sensitive to the extracellular Ca^2+^ as the addition of A23187, an ionophore causing Ca^2+^ leakage from BC, abolishes the BC Ca^2+^ cycles. D) shows the BC Ca^2+^ cycles in correlation with the BC contractions are maintained independent of the ER Ca^2+^. E) A23187 results in one massive BC contraction and the BC Ca^2+^ drop are very closely correlated F) BC does not expand again after massive contraction. Scale Bar B and C) 5 µm F) 10 µm.

### BC Ca^2+^ drop triggers BC contraction

We imaged the BC (30 sec intervals) in the presence of Fluo-8 to test for association between contractions and BC Ca^2+^ levels. Fluctuations in BC Ca^2+^ intensity were observed to correlate with contractions (Figure 2A-D and supporting figure 1E). Kymograph have been plotted using center lines of the BC. Each group of continuous signals represents one contraction cycle (Figure 2C and Supporting figure 1F). The signal from Fluo-8 bleached after 40 min, so we restricted our imaging to 40 min (Black box in Supporting figure 1E). Specifically, the Ca^2+^ levels in BC dropped upon contraction (Supporting Video 3). The rate of Ca^2+^ drop was much faster than the rate of BC contractions. However, the 30 sec imaging intervals obscured the detailed temporal relationship between BC contraction and BC Ca^2+^ depletion. Slow frame-rate was required for 40 min duration, so to observe finer-scale dynamics, we imaged BC at ∽2 sec intervals (Figure 2E) for 5-10 minutes each. Videos containing any contraction were quantified. Results show the BC Ca^2+^ drop started prior to BC contraction (Figure 2F), with a time difference of ∽10 seconds (10.37±1.17 sec) between the start of the Ca^2+^ drop and the start of the BC contraction (Figure 2F and supporting figure 2A-B). This indicates that BC contraction is not the cause of BC Ca^2+^ removal, and leaves open the possibility that BC Ca^2+^ removal might be a cause of BC contraction. We could detect a small increase in Ca^2+^ inside the surrounding hepatocytes near the BC interface, at the same time as removal of Ca^2+^ from BC (Supporting figure 3 C and D). However, Fluo-8 is not ideal for detecting Ca^2+^ in hepatocytes due to low ratio of signal to laser power, cytotoxicity, and photobleaching (Supporting figure 3 E and F). For intracellular imaging we used the Ca^2+^ sensor GCaMP^20^.

### Ca^2+^ waves in hepatocytes are associated with BC contractions

We transfected the Ca^2+^ sensor GCaMP into hepatocytes (Figure 3A). The most optimized transfection protocol has transfection efficiency of ∽1% for primary rat hepatocytes. In the hepatocytes that were transfected, we detected transient increases in Ca^2+^ near BC, followed by apico-basal Ca^2+^ waves (Figure 3B and C). However, the maximum Ca^2+^ intensity decreased as it travelled through cell; and these waves could only be tracked with high laser power (Supporting Figure 4A). While most Ca^2+^ waves originated at the BC interface, we also observed some Ca^2+^ waves originated from cell-cell contacts (R4 in Figure 3B). We presume these non-BC Ca^2+^ waves originated from Ca^2+^ transfer across gap junctions. The non-BC waves showed no temporal association with BC contractions, but when we observed the BC lumen, each BC contraction was associated with a transient increase in intracellular Ca^2+^ at the BC interface (Figure 3C and D, supporting figure 3 B and C, and supporting video 4). In considering cells that had more than 1 BC (Figure 3A), a Ca^2+^ wave that started at a particular BC interface could travel through the cell to another BC interface, but the intensity was much reduced and did not lead to a second contraction at the distal BC. For instance, the Ca^2+^ wave that originated at R2 (large red arrowhead in Figure 3B) also travelled to R1 and R3 (large red arrowhead) but its intensity decreased during transit, and was not associated with BC contractions at R1 or R3 (Figure 3D).

The contractions can be expressed mathematically as a decrease in the slope of BC area over time (Figure 3E); and we found a significant decrease in slope after the Ca^2+^ wave (Figure 3F). Our studies of spatiotemporal correlation are entirely consistent with the hypothesis that BC contractions are caused by Ca^2+^ transfer from BC to surrounding hepatocytes, but the observations are not consistent with hypothetical mechanisms of contractile calcium signaling from gap junctions or cross-cell propagation.

### BC contracts independently from the Ca^2+^ release from ER

ER is an important Ca^2+^ store in cells. It has been proposed previously^15^ that apico-basal Ca^2+^ waves in hepatocytes might be due to the Ca^2+^ release from ER (which surrounds BC) and IP3 generation in response to growth factors ^14^. We imaged BC in the presence of multiple ER inhibitors (Figure 4A) such as 1 µM Thapsigargin (Figure 4B and D and Supporting Figure 4) ^21, 22^, 5 µM Cyclopiazonic acid ^23, 24^ and 1 µM 2,5-di-(ter-butyl)-1,4-benzohydroquinone (tbuHBQ) (Supporting figure 5)^25^. Thapsigargin is a specific and potent inhibitor of sarco/endoplasmic reticulum Ca^2+^-ATPase (SERCA). It forms a dead-end complex and depletes ER Ca^2+^. Cyclopiazonic acid competes with the ATP binding site of SERCA and leads to ER Ca^2+^ depletion, whereas tbuHBQ blocks receptor-activated Ca^2+^ entry, leading to ER leakage^26^. BC contractions still occurred in the presence of any of these drugs, with no change in the Ca^2+^ cycles, nor their temporal association with BC contractions. This indicates that BC contractions are independent of ER Ca^2+^ stores and SERCA pumps. Similar results were obtained in the presence of 3DuM Xestospongin C (Supporting figure 5) ^27, 28^, a reversible IP3 receptor antagonist^29^. These results show that BC contractions do occur independently of IP3-dependent activation of ER Ca^2+^ release.

**Figure 5:**
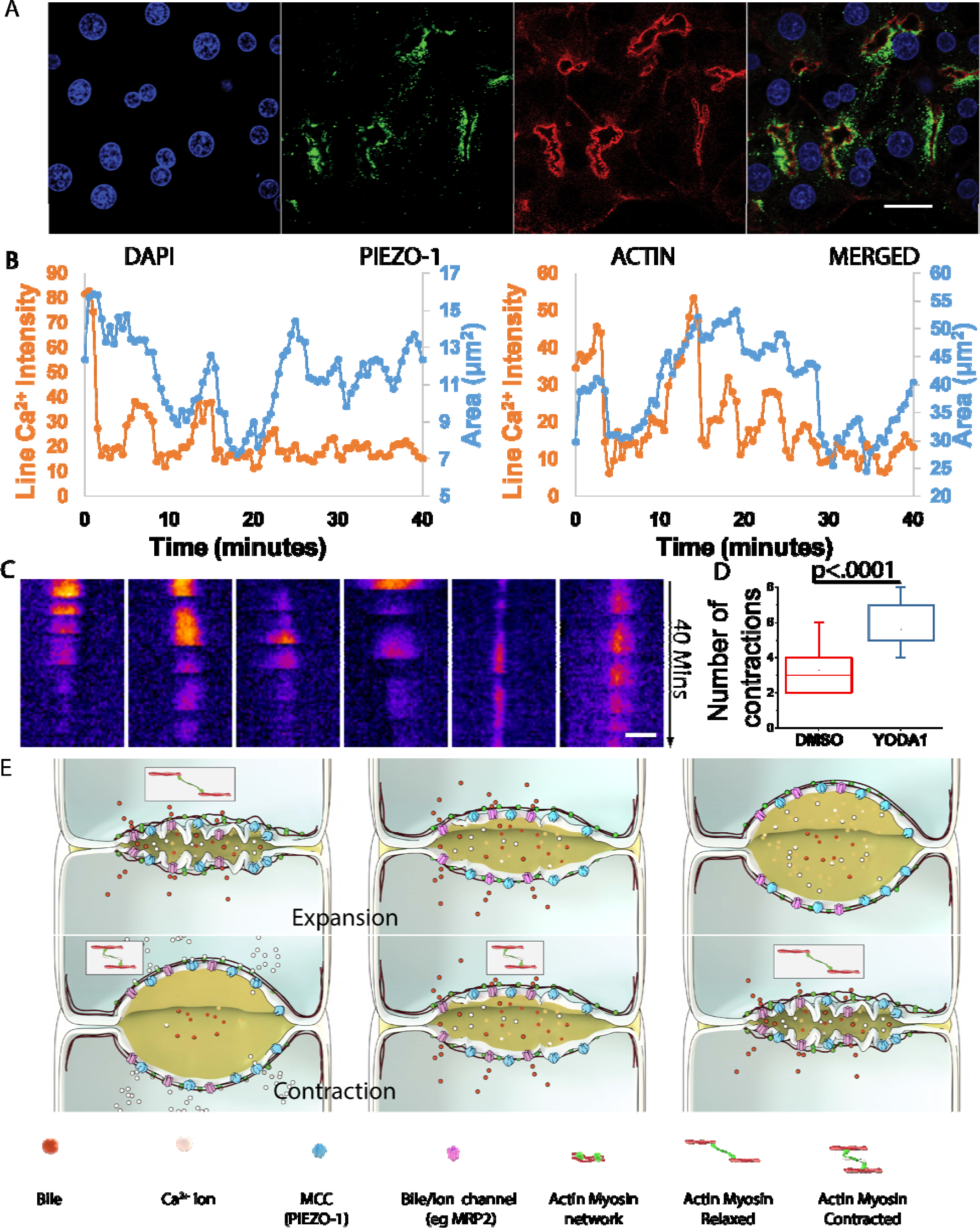
Mechanosensitive Ca^2+^ channel Piezo1 is involved in the BC Ca^2+^ mediated BC contraction. (A) We investigated the localization of various MCC in hepatocytes and found Piezo1 to be localised around BC. To verify if Piezo1 is the major MCC regulating BC contractions, we used 10 µM Yoda-1, an activator of Piezo1 that reduces the pressure threshold to open. (B) In the presence of Yoda-1, we observed a rapid decrease in Ca^2+^ intensity within the first 10 minutes suggesting opeining of Piezo1 resulting in the lower peak Ca^2+^ intensity for subsequent contractions. C) This can be demonstrated in the kymographs (from 3 independent experiments). D) Yoda-1 also led to more frequent BC contraction with less strength. E) Based on these result we propose a model for the BC Ca^2+^ mediated BC contractility. BC membrane has multiple transporter proteins (such as MRP2). These transporter protein constantly secrete content into BC leading to BC expansion and increased intra-canalicular pressure. As pressure threshold for Piezo1 is reached it opens up leading to Ca^2+^ efflux from BC to cells. This Ca^2+^ transfer is seen as the apico-basal Ca^2+^ wave and Ca^2+^ interacts with actomyosin cortex surrounding BC to contract it in cyclic manners. In the presence of Yoda-1, pressure threshold for Piezo1 activation is reduced, as pressure inside BC increases it opens at lower pressure leading to increased BC contraction frequency with less strength. Scale Bar A) 20 µm. C)5 µm

In the presence of 2 μM A23187, a Ca^2+^ ionophore (Supporting figure 4) ^9^, ^30^, we observed that BC Ca^2+^ intensity dropped, followed by BC contraction and collapse, within 10 minutes (Figure 4 C, E and F). A23187 removes the Ca^2+^ gradient across the apical membrane (the BC interface); and in the absence of this gradient, BC never recovered (Figure 4F). This destruction of BC morphology by the A23187 ionophore further supports the hypothesis that Ca^2+^ shuttling between BC and hepatocytes determines BC morphology and cytoskeletal mechanics around the BC cortex.

### Piezo-1 acts as a tension sensor in BC

We hypothesized that increased intra-canalicular pressure (ICP) could activate the opening of mechanosensitive calcium channel (MCC) to allow Ca^2+^ transport at the BC membrane. We tracked the subcellular localization of four MCCs (Supporting figure 6)^31^ and found that Piezo1 is localized at the BC membrane (Figure 5A and supporting video 5). Piezo1 is a calcium channel that is regulated to open whenever membrane pressure exceeds a threshold level. To confirm the role of ICP in regulating Piezo1 activity, we used 10 µM Yoda-1, an activator of Piezo1 that reduces the pressure threshold for Piezo1 to open. In the presence of Yoda-1, we found a significant increase in the frequency of BC contractility (Figure 5B-D), indicating more frequent opening of the MCC is sufficient to cause more frequent contraction of BC. Furthermore, the maximum Ca^2+^ intensity observed during these BC contractions was lower, suggesting there may be a decreased Ca^2+^ gradient across the BC membrane and/or lower ICP as a result of Yoda1-induced sensitization.

**Figure 6:**
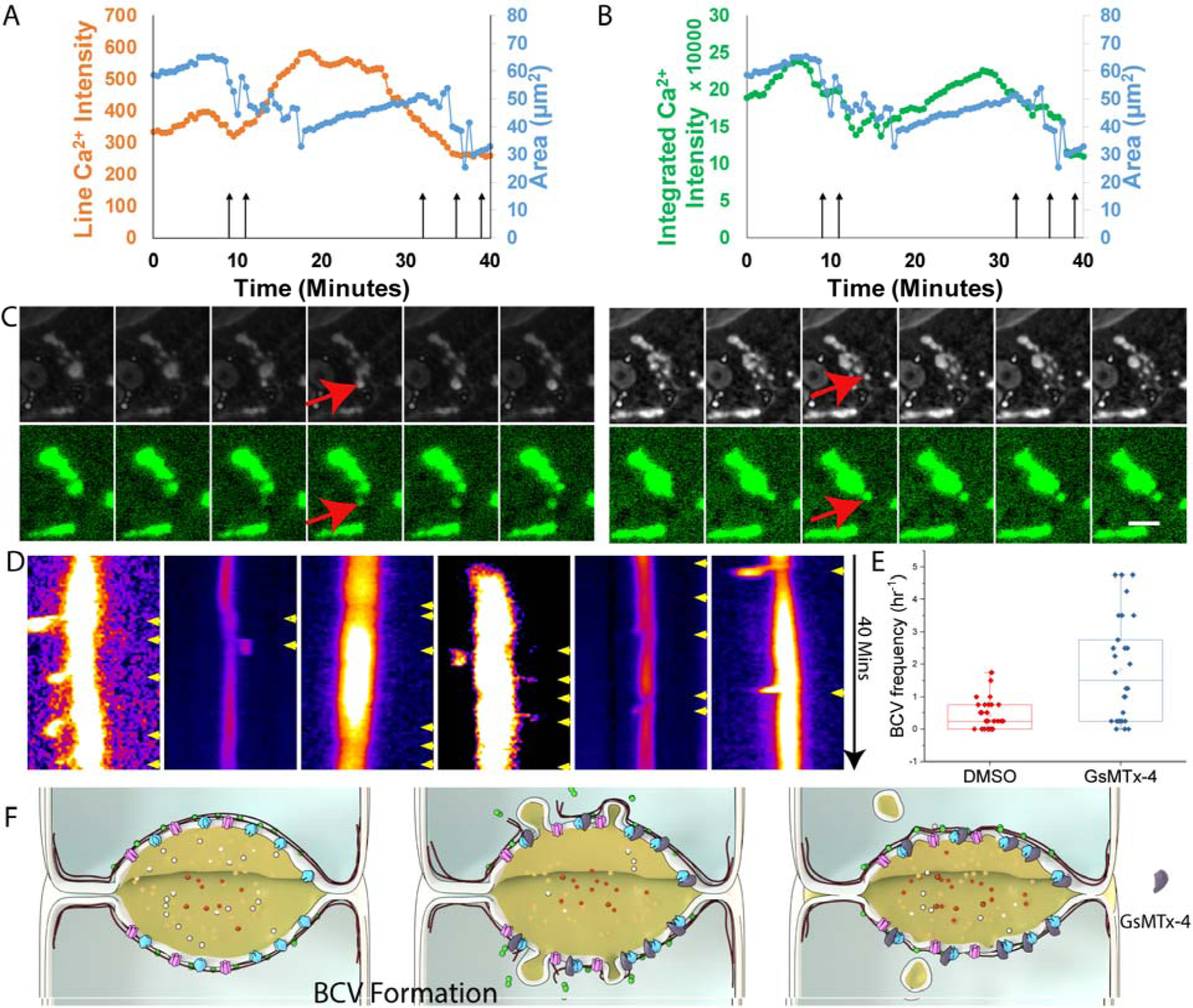
Mechanosensitive Ca^2+^ channel Piezo1 regulates BC contraction and inhibition using GsMTx-4 induces BCV driven contraction. (A) To confirm if Piezo1 regulates BC contraction, we used 10 µM GsMtx-4, an inhibitor of mechanosensitive Ca^2+^ channels. GsMtx-4 abolished the highly synchronized BC Ca^2+^ level and BC contractions. We did not observe any sharp decrease in BC Ca^2+^ intensity. Total Ca^2+^ signal is still correlated with the BC area. In the presence of GsMTx-4, BC area decreases and the total Ca^2+^ signal reduced due to the BCV formation (black arrow in A and B) as shown in C (scale bar 10 µm). D) This can be seen in all the kymographs (from 3 independent experiments) wherein no sharp Ca^2+^ decrease is observed. Yellow arrowheads marks the BCV. E) shows quantification of BCV frequency (n=30). The BCV frequency is s8 fold higher in the presence of GsMTx-4. F) Illustrates a model showing that GsMTx-4 blocks MCC leading to increased pressure and BCV formation.

We next inhibited the mechano-activation of Piezo1 (the opposite intervention) using 10 μM GsMTx-4 Piezo^32^. In the presence of GsMTx-4, we observed no instances of Ca^2+^ depletion from the BC and no BC contraction cycles (Figure 6A-D, flat kymograph), while the control kymograph (Figure 2D) showed periodic BC Ca^2+^ cycles. We could no longer find a correlation between mean Ca^2+^ (measured as “line Ca^2+^ intensity”) and BC area, indicating that GsMTx-4 was sufficient to abrogate calcium transport at the BC interface and to halt BC contractions. This suggests that MCC activity is required for calcium transport at the BC interface and is required for BC contractions. There is a correlation between total BC area and total Ca^2+^ Figure 6B, but this may be a trivial finding because bile is rich in calcium and any non-contractile changes in BC area, such as pinching off BCV, must result in simultaneous changes in total BC volume and total BC calcium (Figure 6 C). Inability to transfer BC Ca^2+^ to cells halted contraction, which is presumed to impair the propulsion of contents and the release of ICP. In such cases, cells resort to remodeling through BCV, which resembles to some extent the cytoskeletal mechanics of blebbing^33^. The presence of GsMTx-4 caused ∽8-fold higher frequency of BCV (Figure 6E), leading to a complete suppression of BCV-independent contractions, which is a recognized BC response to cholestasis^33^. The agonist and inhibitor of Piezo1 show that Piezo1 is required for transfer of Ca^2+^ from BC to surrounding hepatocytes and is required for cyclic BC contractions (Figure 5E and 6F).

## Discussion

Cells and tissues can sense and respond to changes in their local mechanical environment^34^. For instance, blood vessel contractility and RBC size are altered in response to changes in blood flow; and mechanosensitive channels (MCC) such as Piezo play a central role in pressure sensing to regulate blood flow^35-37^. Analogous regulation occurs in esophageal food sensing and the initiation of peristalsis^38^. In these and various other lumenal structures, the volume of contents can in itself regulate the contractility of tissues by stimulating muscle cells^39^. There are other lumenal structures in the body where contractility does not depend on the activity of muscle cells^40^. Here we have investigated the contractility in one such lumenal structure, BC, and found that local pressure sensing via MCC play a role in regulating contractility.

Physiologically the contractile nature of BC is supposed to propel bile from BC to bile duct^2, 3, 7, 9, 13, 41^. However, the mechanism of the contraction process is poorly understood. Previous studies from ourselves and others suggested that the BC contractility is a cyclic process and each cycle consists of at least two phases^9, 10, 13, 19, 33, 42^. The first phase is expansion, wherein the BC expands due to activity of transporter proteins such as MRP2, BSEP and others^2, 43-45^. In the second phase, following expansion, the lumen contracts. These contractions are important for volume reduction because the lumen membrane is finite and cannot tolerate limitless BC expansion. In order to contract the BC, the actomyosin network surrounding the lumen contracts^33^. A BC can reduce its volume or contract *in vitro* by two different mechanisms. One method is by para-cellular leak^42^ and other is by the formation of BCV ^33^. In case of increased pressure^46^ and compromised cytoskeleton, BC favors BCV-driven mechanisms^33^.

Growth factors and hormones such as vasopressin trigger the release of IP3 in cells. Released IP3 then binds to IP3R on the ER membrane, causing the ER to release Ca^2+^ and this Ca^2+^ has been postulated to cause BC contractions^12^. Intracellular Ca^2+^ can cause contractions; however, whether growth factor-induced Ca^2+^ release from the ER causes BC contractions is not proven. There are reports suggesting an increase in intracellular Ca^2+^ in hepatocytes in the absence of IP3 generation. Other studies have suggested a role of extracellular Ca^2+^ rather than ER Ca^2+^ in regulating BC contraction^16, 47^. Even in the absence of any growth factor, BC expand and contract periodically *in vitro*. Thus, we hypothesize that the trigger and control unit for BC contractions might not be extra-canalicular but from BC itself. To test this hypothesis, we used Ca^2+^ dyes to image Ca^2+^ and found that Ca^2+^ shuttles inside and outside of BC; and Ca^2+^ depletion from BC precedes BC contraction.

Using GCaMP for intracellular Ca^2+^ measurements, we found that most Ca^2+^ waves originated from the BC membrane. BC-originating Ca^2+^ waves were associated with a significant decrease in BC area, signifying a tight temporal association with BC contractions. This is in contrast to previous reports that the BC membrane was associated with the ER which releases Ca^2+^ under IP3 regulation. In order to test whether BC contractions depend on ER or IP3 generation in cells, we used multiple SERCA inhibitors and Xestospongin C, a competitive inhibitor of IP3. None of these inhibitors could stall BC contractions or change the BC Ca^2+^ cycles. BC contractions and BC Ca^2+^ cycles were inhibited by A23187, an ionophore, supporting that extracellular Ca^2+^ (e.g. BC) instead of ER Ca^2+^ regulate BC contractions. We further screened MCC and found that Piezo-1 is localized on the BC membrane. Currently there are a few classes of ion channels such as Piezo-1 that are considered truly mechanosensitive^48^. Piezo-1, a cationic MCC, responds to lateral membrane tension^49^ and can sense mechanical forces in membrane^48^. As BC expands, the BC membrane tension increases beyond certain threshold; Piezo-1 should open causing Ca^2+^ leakage into the cells surrounding BC (Figure 5E). To test this hypothesis, we treated hepatocytes with Yoda1, a specific potent activator of Piezo1^50^. Yoda1 decreases the pressure threshold for Piezo1 activation which should increase BC contractility. We observed approximately two-fold increase in BC contractility confirming the role of Piezo1 in regulating Ca^2+^ transfer from BC and BC contractions. Further, we used GsMTX-4 peptide, the most widely used Piezo-1 Inhibitor^32^ ^51^. We observed a complete suppression of BC contraction. Ca^2+^ was retained in BC and not removed nor show cyclic fluctuations. We also observed BCV-driven contractions with eight-fold increase in the BCV formation indicating activation of cholestatic response. Thus, BC is an autonomous contractile entity and its contractility is independent of external factors such as hormones or growth factor.

MCC are required for multiple functions such as hearing and substrate stiffness sensing^52^. In addition, MCC have been shown to play a major role in regulating endothelial lumen dynamics by sensing intra-vascular pressure^35, 36, 53^. However, the role of MCC in regulating the pressure in secretory lumens (such as acini, canaliculi, blastocyst) is not clear. Here we have shown an example of a secretory lumen, BC, where the MCC Piezo1 regulated the contractile dynamics, suggesting a feedback mechanism for secretion-driven pressure to activate propulsion and homeostasis. Identification of Piezo-1 as a molecular target provides an opportunity to further develop therapeutic agents that might provide an alternate therapy for cholestatic conditions.

## Acknowledgements

We acknowledge funding support from the Mechanobiology Institute (National University of Singapore, Singapore, Grant number: R-714-006-008-271), Institute of Bioengineering and Nanotechnology, Biomedical Research Council IAF-PP (A*STAR, Singapore), NUHS Innovation Seed Grant 2017 (R-185-000-343-733); Singapore-MIT Alliance for Research and Technology, National Medical Research Council (Ministry of Health, Singapore, Grant Number: R-185-000-294- 511) and MOE ARC (R-185-000-342-112) to HY. LTK and BPN were supported by the Singapore Ministry of Health’s National Medical Research Council (NMRC) under its Open Fund Individual Research Grant scheme (OFIRG15nov062).

Diego Pitta de Araujo, Animator, Science Communications Core, Mechanobiology Institute, National University of Singapore designed all the animations. Hui Ting, Image Processing Core, Mechanobiology Institute helped with image processing. We also thank the Microscopy Core at the Mechanobiology Institute, and NUHS core facilities, National University of Singapore for their help with microscopy. KG is a Mechanobiology Institute PhD scholar.

Material and Methods

1. Chemicals: Linoleic acid, 10x phosphate buffer saline (PBS), William’s E medium, bovine serum albumin (BSA), glutamine, dexamethasone, penicillin, insulin, sodium hydroxide, were obtained from Sigma Aldrich, USA. Type I bovine dermal collagen solution was obtained from Advanced Bio Matrix, USA. Calcium Green™-1, AM, Lipofectamine^®^ 2000 Transfection Reagent was purchased from Thermo Scientific, USA. 2,5-di-(ter-butyl)^-1^,4-benzohydroquinone (tbuHBQ) (ab120276), Thapsigargin (ab120286), A23187 (ab120287), Cyclopiazonic acid (ab120300), Xestospongin C (ab120914), Fluo-8 AM were purchased from Abcam, United Kingdom. SiR-Actin Kit containing SiR-Actin and Verapamil was purchased from Cytoskeleton, Inc., USA. GsMTx-4, Anti Peizo-2, Anti-TRPC-4, Anti-TRPC-6 and Anti Peizo-1 Antibody (host: rabbit) were obtained from Almone lab, Israel.
2. Isolation and culture of rat hepatocytes: Hepatocytes were isolated from male Wistar rats (weight 200-300gm) via two step collagenase perfusion method. The animals were obtained from InVivos, Singapore. Animals were handled in accordance to the IACUC protocol approved by the IACUC committee of the National University of Singapore, protocol number R15–0027. ∽300 million cells were isolated with viability above 90% as determined by the Trypan blue exclusion assay.
3. Collagen sandwich culture. Isolated hepatocytes were cultured in collagen sandwich culture system. 35 mm culture dishes were coated with 1.5 mg mL^-1^ neutralized Type I bovine dermal collagen solution diluted in 0.01 M hydrochloride for 2 h. Coated dishes were washed twice with 1x PBS, and 0.8 million cells were seeded in the dish. Hepatocytes were allowed to attach for 2 h. Unattached cells were removed by washing twice in 1x PBS. Hepatocytes were then incubated for an additional 1 h to allow spreading. Following which, a layer of collagen solution was pipetted over the cells (1mL of 0.5 mg mL^-1^ neutralized collagen solution per well). 1.5 mg mL^-1^ neutralized collagen was obtained by mixing 3 mg mL^-1^ of acidic collagen with a neutralizing solution (in equal parts). The neutralizing solution comprises 0.1M sodium hydroxide, 10x PBS, 1x PBS in a volume ratio of 1:1:6 respectively. Working concentration of the neutralized collagen (0.5 mg mL^-1^) was then obtained by diluting the 1.5 mg mL^-1^ neutralized collagen solution with culture medium. Culture medium comprises William’s E medium supplemented with 2 mM L-Glutamine, 1 mg mL^-1^ BSA, 0.3 μg mL^-1^ of insulin, 100 nM dexamethasone, 50 μg mL^-1^ linoleic acid, 100 unit mL^-1^ penicillin, and mg mL^-1^ streptomycin. Cells were incubated with 5% CO2 at 37°C and 95% humidity. Culture medium was changed daily.
4. Labelling of hepatocytes with SiR-Actin: Hepatocytes were labelled with SiR-Actin using manufacturer protocol. Briefly, 40 hours after seeding hepatocytes, Hepatocyte media was changed to 1mL fresh WE media (containing 200nM SiR actin). Hepatocytes were incubated for 1h and then imaged using an inverted confocal microscope (Nikon A1R). 1 μM Verapamil was added for taking static images but was not added in samples used for time lapse images. Verapamil is a transporter inhibitor. It prevents SiR-Actin from going into BC and gives a clearer image, however we found that it can also inhibit contractility thus it was not added in samples used for tracking BC. In absence of Verapamil some SiR-Actin did go to BC, however the contrast was still high to segment BC boundary.
5. Transfection of hepatocytes: Hepatocytes were transfected with GCaMP using Lipofectamine^®^ 2000 according to the manufacturer’s protocol but with modifications for use with primary rat hepatocytes. DNA:Lipofectamine ratio and amount were optimized based on transfection efficiency and viability. Under optimized conditions, the transfection mixture was prepared by adding 2 μg of the plasmid dissolved in 250 μL of antibiotic-free William’s E medium to a solution containing 4 μL of Lipofectamine dissolved in 250 μL of antibiotic-free William’s E medium. This solution was the incubated at room temperature for 20 min and then added to hepatocytes cultured on collagen-coated glass 35-mm dishes (IWAKI) for 4 h. Hepatocytes were incubated with the transfection mixture overnight. Hepatocytes were then washed and overlaid with collagen as described earlier for sandwich culture. Under these optimized transfection conditions, transfection efficiency of approximately 2% was achieved within 24 h of transfection. Plasmids used in this study were expanded in E. coli (DH5α™ competent cells, ThermoFisher), and extracted using the Qiagen Maxiprep kit following the manufacturer’s protocol. The dynamics of Ca^2+^ in labelled cells was monitored using an inverted confocal microscope (Nikon A1R). Time-lapse images were captured at time intervals of 1-2 s (or fastest possible) for 5-10 minute. Images were analyzed as described in Imaging Analysis.
6. Labelling Hepatocytes with Ca^2+^ dyes: We used multiple Ca^2+^ dyes to image Ca^2+^ in hepatocytes as per manufacturer protocol. Fluo-8 was primarily used in all the experiments as it was most stable for long term imaging. 40 hours after seeding hepatocytes, Hepatocyte media was changed to 1mL fresh WE media (containing 4 µM Fluo-8) and incubated for 30 minutes. Following incubation, the cells were imaged at 30 sec intervals for 40 minutes using Nikon IMQ at 20X magnification. For faster imaging, hepatocytes were also stained with SiR actin and imaged using inverted confocal microscope (Nikon A1R) at 1-2 second interval for 5-10 minutes. We saw cell death after longer imaging time and thus restricted faster imaging to less than 10 minutes.
7. Drug treatment of hepatocytes: 40 hours after seeding hepatocytes, Hepatocyte media was changed to 1mL fresh WE media (containing 4 µM Fluo-8) and incubated for 30 minutes. Following incubation, 1mL fresh WE media containing drug (either tbuHBQ, Thapsigargin, A23187, Cyclopiazonic acid, Xestospongin C, glyburide, verapamil or GsMTx-4) was added to give desired drug concentration. The drugs were added while sample was placed on microscope and imaged immediately after drug addition. Cells were imaged at 30 sec intervals for 40 minutes using Nikon IMQ at 20X magnification.
8. Image processing: For long term images X-Y drift (in any) was corrected in IMARIS and corrected images were analyses either using MARLAB, Fiji or IMARIS. Phase contrast images obtained using Nikon biostation IMQ were segmented using a custom made software. We develop a software for segmentation of a bile canaliculus from a time-lapse phase-contrast image. The software is written in MATLAB and is based on a combination of interactive thresholding, morphological image processing, pattern matching, and manual segmentation. A user friendly graphical user interface (GUI) is also implemented. At the beginning, a region of interest (ROI) for a bile canaliculi to be segemented is selected. In the software software, each image frame is associated with the following information: 1) a threshold for thresholding on the frame, 2) boundary1 storing the canaliculus boundary by automatic thresholding, 3) boundary2 storing the canaliculus boundary by manual segmentation or by manual selection (if any), 4) status1 (status1 > 0 indicates that there is a manual region selection/segmentation/merging in the frame), and 5) status2 (status2 > 0 indicates that the frame is segmented automatically based on the segmentation from the previous frames). Before the image frames are processed, all the intensities are normalized to the range of [0, 1], and a default threshold value within that range is set for all the frames. Also, boundary1 and boundary2 are set as empty, and status1 = status2 = 0 for each frame. At the beginning, the current frame is set as the first frame. Then thresholding with the default threshold is applied on the first frame. Segmented regions are displayed on the screen, and the largest region is selected for boundary1. For each image frame, the user can choose to execute the operations such as changing threshold, selecting correct ROI, merging to ROIs, segmenting BC manually, Automatic processing all the remaining frames and saving the result. BCV frequency was manually calculated by visualizing individual BC in Fiji. For determination of Ca^2+^ waves, a small ROI was drawn at BC surface and intensity measured across time lapse images. BC area form SiR actin stained images were auto thresholded in Fiji and area measured using in built plugin.
9. Image Analysis: Ca^2+^ level fluctuation inside BC were represented as kymographs. Kymograph are drawn along a line drawn in middle of BC. Most of the kymograph show a time scale of 40 minutes unless otherwise stated. We have to limit imaging to 40 minutes as at longer imaging time we had issues with bleaching (supporting figure 1F). Also Ca^2+^ intensity during the contraction is measured using 1) Integrated intensity for the area equivalent to maximum projected area for all the frames captured and is mentioned as “Integrated Ca^2+^ Intensity” 2) a line scan across kymograph and is mentioned as “Line Ca^2+^ intensity”. First methods describe the total Ca^2+^ in the BC and is sensitive to both area change and Ca^2+^ level drop in BC whereas second method is sensitive only to Ca^2+^ level drop in BC (fast removal of Ca^2+^ from BC) (supporting figure 1F-I).
10. Antibody staining: Hepatocytes cultured for 40 hours were fixed with 4% paraformaldehyde (PFA) for 20 min at 37 °C and permeabilized thereafter with 0.1% Triton X-100 in PBS for 10 min. Nonspecific binding was prevented by blocking the permeabilized cells for 1 hour in a PBS solution containing 5% BSA at room temperature. Cells were then incubated with primary antibodies for PEIZO-1, PEIZO-2 TRPC-6 or TRPC-4 (1:100) in 1X PBS solution with 5% BSA overnight at 4 °C. Cells were then washed five times with gentle shaking in 1X PBS and incubated for 2 hours with rabbit mouse secondary antibodies (1:100) at room temperature. After washing five times in PBS, nuclei were stained for 30 min with the nuclear dye DAPI (1:1,000) and actin was stained with phalloidin (1:200). Cells were then washed five times in 1X PBS and imaged using Nikon A1R. Images were processed using IMARIS (Bitplane Technologies, USA) and fiji.
11. Statistical Analysis: Data values in this report are presented as average ± standard error of the mean (SEM). In most groups, the initial test of difference across groups was analyzed using one-way ANOVA. The Student’s t test was then used as a post-hoc test to identify the significant differences among the different conditions in the experiment. p values <0.05 (∗), p <0.01 (∗∗) were considered statistically significant. For testing significance of change in slop with Ca^2+^ wave, sign test was also used to analyse the significance of the reduction of BC area.

## Notes

Conflict of interest: Dr. Yu reports personal fees and non-financial support from Invitrocue, Histoindex, Pishon Biomedical Co. Ltd., outside the submitted work. All other authors have nothing to disclose.

## References

1. Li, Q. et al. Extracellular matrix scaffolding guides lumen elongation by inducing anisotropic intercellular mechanical tension. Nat Cell Biol 18, 311–318 (2016).

2. Wagner, M., Zollner, G. & Trauner, M. New molecular insights into the mechanisms of cholestasis. J Hepatol 51, 565–580 (2009).

3. Watanabe, N., Tsukada, N., Smith, C.R. & Phillips, M.J. Motility of bile canaliculi in the living animal: implications for bile flow. The Journal of cell biology 113, 1069–1080 (1991).

4. LeCluyse, E.L., Fix, J.A., Audus, K.L. & Hochman, J.H. Regeneration and maintenance of bile canalicular networks in collagen-sandwiched hepatocytes. Toxicology in Vitro 14, 117–132 (2000).

5. Burbank, M.G. et al. Early Alterations of Bile Canaliculi Dynamics and the Rho Kinase/Myosin Light Chain Kinase Pathway Are Characteristics of Drug-Induced Intrahepatic Cholestasis. Drug Metab Dispos 44, 1780–1793 (2016).

6. Reif, R. et al. Bile canalicular dynamics in hepatocyte sandwich cultures. Arch Toxicol 89, 1861–1870 (2015).

7. Arias, I.M. et al. The biology of the bile canaliculus, 1993. Hepatology (Baltimore, Md.) 17, 318–329 (1993).

8. Sudo, R., Kohara, H., Mitaka, T., Ikeda, M. & Tanishita, K. Coordinated movement of bile canalicular networks reconstructed by rat small hepatocytes. Annals of biomedical engineering 33, 696–708 (2005).

9. Sumio, W. et al. Bile canalicular contraction in the isolated hepatocyte doublet is related to an increase in cytosolic free calcium ion concentration. Liver 8, 178–183 (1988).

10. Watanabe, S. & Phillips, M.J. Ca2+ causes active contraction of bile canaliculi: direct evidence from microinjection studies. Proceedings of the National Academy of Sciences of the United States of America 81, 6164–6168 (1984).

11. Tsuneo, K., Ulrike, B., Zenaida, G. & M., A.I. Extracellular ATP, intracellular calcium and canalicular contraction in rat hepatocyte doublets. Hepatology (Baltimore, Md.) 14, 640–647 (1991).

12. Combettes, L., Dumont, M., Berthon, B., Erlinger, S. & Claret, M. Release of calcium from the endoplasmic reticulum by bile acids in rat liver cells. The Journal of biological chemistry 263, 2299–2303 (1988).

13. Yokomori, H. et al. Bile canalicular contraction and dilatation in primary culture of rat hepatocytes--possible involvement of two different types of plasma membrane Ca(2+)-Mg(2+)-ATPase and Ca(2+)-pump-ATPase. Medical electron microscopy: official journal of the Clinical Electron Microscopy Society of Japan 34, 115–122 (2001).

14. Amaya, M.J. & Nathanson, M.H. Calcium signaling in the liver. Comprehensive Physiology 3, 515–539 (2013).

15. Nagata, J. et al. Lipid rafts establish calcium waves in hepatocytes. Gastroenterology 133, 256–267 (2007).

16. Sharanek, A. et al. Rho-kinase/myosin light chain kinase pathway plays a key role in the impairment of bile canaliculi dynamics induced by cholestatic drugs. Scientific Reports 6, 24709 (2016).

17. Seglen, P.O. Preparation of isolated rat liver cells. Methods in cell biology 13, 29–83 (1976).

18. Gupta, K. et al. 6.28 Liver Tissue Engineerings A2 - Ducheyne, Paul, in Comprehensive Biomaterials II 491–512 (Elsevier, Oxford; 2017).

19. Luo, X. et al. Directed Differentiation of Adult Liver Derived Mesenchymal Like Stem Cells into Functional Hepatocytes. Scientific Reports 8, 2818 (2018).

20. Akerboom, J. et al. Optimization of a GCaMP Calcium Indicator for Neural Activity Imaging. The Journal of Neuroscience 32, 13819–13840 (2012).

21. Wictome, M., Henderson, I., Lee, A.G. & East, J.M. Mechanism of inhibition of the calcium pump of sarcoplasmic reticulum by thapsigargin. Biochemical Journal 283, 525–529 (1992).

22. Oslowski, CM. & Urano, F. Chapter Four - Measuring ER Stress and the Unfolded Protein Response Using Mammalian Tissue Culture System, in Methods in Enzymology, Vol. 490. (ed. P.M. Conn) 71–92 (Academic Press, 2011).

23. Grigoryan, G. & Segal, M. Prenatal stress alters noradrenergic modulation of LTP in hippocampal slices. Journal of neurophysiology 110, 279–285 (2013).

24. Komori, Y. et al. Ca2+ homeostasis, Ca2+ signalling and somatodendritic vasopressin release in adult rat supraoptic nucleus neurones. Cell Calcium 48, 324–332 (2010).

25. Krause, E., Pfeiffer, F., Schmid, A. & Schulz, I. Depletion of Intracellular Calcium Stores Activates a Calcium Conducting Nonselective Cation Current in Mouse Pancreatic Acinar Cells. Journal of Biological Chemistry 271, 32523–32528 (1996).

26. Nelson, E.J. et al. Inhibition of L-type calcium-channel activity by thapsigargin and 2,5-t-butylhydroquinone, but not by cyclopiazonic acid. The Biochemical journal 302 (Pt 1), 147–154 (1994).

27. Miyamoto, S. et al. Xestospongin C, a selective and membrane-permeable inhibitor of IP(3) receptor, attenuates the positive inotropic effect of α-adrenergic stimulation in guinea-pig papillary muscle. British Journal of Pharmacology 130, 650–654 (2000).

28. Gafni, J. et al. Xestospongins: potent membrane permeable blockers of the inositol 1,4,5-trisphosphate receptor. Neuron 19, 723–733 (1997).

29. Ozaki, H. et al. Inhibitory mechanism of xestospongin-C on contraction and ion channels in the intestinal smooth muscle. Br J Pharmacol 137, 1207–1212 (2002).

30. Przygodzki, T., Sokal, A. & Bryszewska, M. Calcium ionophore A23187 action on cardiac myocytes is accompanied by enhanced production of reactive oxygen species. Biochimica et Biophysica Acta (BBA) - Molecular Basis of Disease 1740, 481–488 (2005).

31. Martinac, B. Mechanosensitive ion channels: molecules of mechanotransduction. Journal of Cell Science 117, 2449–2460 (2004).

32. Bae, C., Sachs, F. & Gottlieb, P.A. The mechanosensitive ion channel Piezo1 is inhibited by the peptide GsMTx4. Biochemistry 50, 6295–6300 (2011).

33. Gupta, K. et al. Actomyosin contractility drives bile regurgitation as an early response during obstructive cholestasis. J Hepatol 66, 1231–1240 (2017).

34. Yusko, E.C., Asbury, C.L. & Bement, W. Force is a signal that cells cannot ignore. Molecular Biology of the Cell 25, 3717–3725 (2014).

35. Ranade, S.S. et al. Piezo1, a mechanically activated ion channel, is required for vascular development in mice. Proceedings of the National Academy of Sciences 111, 10347–10352 (2014).

36. Rode, B. et al. Piezo1 channels sense whole body physical activity to reset cardiovascular homeostasis and enhance performance. Nature Communications 8, 350 (2017).

37. Cahalan, S.M. et al. Piezo1 links mechanical forces to red blood cell volume. eLife 4 (2015).

38. Lin, Z. et al. The four phases of esophageal bolus transit defined by high-resolution impedance manometry and fluoroscopy. American journal of physiology. Gastrointestinal and liver physiology 307, G437–444 (2014).

39. Goyal, R.K. & Chaudhury, A. Physiology of normal esophageal motility. Journal of clinical gastroenterology 42, 610–619 (2008).

40. Oshio, C. & Phillips, M.J. Contractility of bile canaliculi: implications for liver function. Science (New York, N.Y.) 212, 1041–1042 (1981).

41. Strange, R.C. Hepatic bile flow. Physiological reviews 64, 1055–1102 (1984).

42. Dasgupta, S., Gupta, K., Zhang, Y., Viasnoff, V. & Prost, J. Physics of lumen growth. Proceedings of the National Academy of Sciences of the United States of America 115, E4751–e4757 (2018).

43. Sohail, M.I. et al. Molecular Mechanism of Taurocholate Transport by the Bile Salt Export Pump, an ABC Transporter Associated with Intrahepatic Cholestasis. Molecular pharmacology 92, 401–413 (2017).

44. Trauner, M. & Boyer, J.L. Bile salt transporters: molecular characterization, function, and regulation. Physiological reviews 83, 633–671 (2003).

45. Stieger, B., Fattinger, K., Madon, J., Kullak-Ublick, G.A. & Meier, P.J. Drug- and estrogen-induced cholestasis through inhibition of the hepatocellular bile salt export pump (Bsep) of rat liver. Gastroenterology 118, 422–430 (2000).

46. Cai, S.-y. & Boyer, J.L. Bile Infarcts – new insights into the pathogenesis of obstructive cholestasis. Hepatology (Baltimore, Md.) 0.

47. Bouscarel, B., Fromm, H. & Nussbaum, R. Ursodeoxycholate mobilizes intracellular Ca2+ and activates phosphorylase a in isolated hepatocytes. The American journal of physiology 264, G243–251 (1993).

48. Syeda, R. et al. Piezo1 Channels Are Inherently Mechanosensitive. Cell reports 17, 1739–1746 (2016).

49. Coste, B. et al. Piezo proteins are pore-forming subunits of mechanically activated channels. Nature 483, 176–181 (2012).

50. Syeda, R. et al. Chemical activation of the mechanotransduction channel Piezo1. eLife 4 (2015).

51. Li, C. et al. Piezo1 forms mechanosensitive ion channels in the human MCF-7 breast cancer cell line. Scientific Reports 5, 8364 (2015).

52. Chesler, A.T. & Szczot, M. Portraits of a pressure sensor. eLife 7, e34396 (2018).

53. AbouAlaiwi, W.A. et al. Ciliary polycystin-2 is a mechanosensitive calcium channel involved in nitric oxide signaling cascades. Circulation research 104, 860–869 (2009).

